# Massive expansion of the pig gut virome based on global metagenomic mining

**DOI:** 10.1101/2023.04.22.537307

**Authors:** Jiandui Mi, Xiaoping Jing, Chouxian Ma, Yiwen Yang, Yong Li, Yu Zhang, Ruijun Long, Haixue Zheng

## Abstract

The pig gut virome plays a crucial role in the gut microbial ecosystem of pigs, yet a comprehensive reference database is still lacking. To address this gap, we established the Pig Virome Database (PVD) of the gut that comprises 5,566,804 viral contig sequences from 4,650 publicly available gut metagenomic samples using a pipeline named “metav” developed in this study. The majority of viral operational taxonomic units (vOTUs) were identified as *Caudoviricetes* (65.36%). By clustering sequences, we identified 48,299 vOTU genomes, of which 92.83% were not found in existing major databases. The PVD database contains a total of 18,161,503 protein-coding genes that can be used to explore the functional potential of the pig gut virome. Our study showed that the PVD can improve the detection of viruses that carry antibiotic/metal resistance genes, mobile genetic elements, virulence factor genes, and quorum sensing systems. These findings highlight the extensive diversity of viruses in the pig gut and provide detailed insight into host‒virus interactions.

## Introduction

The gut microbiota is a complex microbial ecosystem that plays crucial roles in pig health, nutrient metabolism, and productivity (Chen et al., 2021a; Hu et al., 2023; Wang et al., 2022; Yang et al., 2022). Pig intestinal virus infections are prevalent in the global pig industry, but the large numbers of diverse viruses that inhabit the pig gut can greatly affect the structure and function of the gut microbial ecosystem (Shkoporov et al., 2022a). Controlling the occurrence of these diseases solely through vaccines or medication is difficult. Thus, understanding the pig gut virome is essential for improving pig health and enhancing production efficiency. However, despite their ubiquity, knowledge of the diversity of the virome in the pig gut ecosystem is limited, and most virome genomes fail to match existing genome databases. Furthermore, most virome databases are established based on human gut metagenomic samples (Camarillo-Guerrero et al., 2021; Gregory et al., 2020; Nayfach et al., 2021b), and there is a lack of a specialized virus database for pigs (Wu et al., 2022a). Therefore, a comprehensive database of the virome from the pig gut microbiota is a prerequisite for characterizing virus diversity, understanding host‒virus interactions, and resolving the functions of viral genomes.

Currently, there is a wealth of metagenomic data available that offers a unique opportunity to discover viral genomes (Edgar et al., 2022). Despite being generated through untargeted methods without virus particle enrichment, these datasets still contain a significant number of viral genomes. Several databases, such as GOV2 (Gregory et al., 2019), IMG/VR4 (Camargo et al., 2023a), GVD (Gregory et al., 2020), MGV (Nayfach et al., 2021b), and GPD (Camarillo-Guerrero et al., 2021), have been established to facilitate the analysis of virome from various environments using metagenomic data. Since the establishment of these databases, the number of publicly available pig gut microbiota datasets has rapidly increased (Chen et al., 2021b; Gaio et al., 2022; Gaio et al., 2021; Gaire et al., 2022; Luiken et al., 2020; Tao et al., 2022; Wu et al., 2022a; Xiao et al., 2016; Zhang et al., 2022).

To make use of the existing resources and provide a comprehensive, global view of the pig gut virome, we developed a comprehensive metav analysis pipeline to examine 4,650 metagenomic samples. We optimized several software tools to improve processing speed and end-to-end output. The Pig Virome Database (PVD) of the gut was established, containing 5,566,804 viral contig sequences estimated to be >50% complete, representing 48,299 vOTU genomes. Our analysis uncovered a diverse and complex pig gut virome, with a high number of unique vOTUs (92.83%) compared to other databases. Moreover, we identified several potential novel viral species and provided evidence of multiple viral species coinfection in the pig gut. These findings enhance our understanding of the pig gut virome, and provide insights into the complexity of gut ecosystems, emphasizing the importance of further research in this field.

## Methods

### Data Assembly and Viral Identification

We developed “metav”, a pipeline for identifying viruses from raw metagenomics generated via next generation sequencing. It comprised three main steps: 1) quality control of raw reads using fastp v.0.23.2 (with parameters --detect_adapter_for_pe and --dont_eval_duplication -w 16), followed by the removal of host (pig genome: Sscrofa11.1) contamination using BWA v.0.7.17 with parameters (-k 31 -p -S -K 200000000) (Chen et al., 2018; Li & Durbin, 2009), 2) assembly of the data using either Megahit v.1.2.9 (with -k-list 39, 59, and 79) or MetaSpades v.3.15.5 (Li et al., 2015; Nurk et al., 2017), and 3) viral identification using VirSorter2 v.2.2.3 and DeepVirFinder v.2.0 (Guo et al., 2021a; Ren et al., 2020). Viral sequences were identified based on VirSorter2’s SOP with (Guo et al., 2021b) and *P*>=0.95 using DeepVriFinder. We collected metagenomics data from various sources, including SRP188615/PRJNA526405 (Gaio et al., 2022; Gaio et al., 2021), CNP0000824 (Chen et al., 2021b), PRJEB11755 (Xiao et al., 2016), PRJNA788462 (Gaire et al., 2022), PRJNA775062(Tao et al., 2022), PRJEB22062 (Luiken et al., 2020), PRJCA009609 (Wu et al., 2022a), PRJEB44118 (Zhang et al., 2022), and 70 samples collected in our lab. In total, 4,650 samples were used in our study to extract the viral contigs. Each sample was processed using the above pipeline, and then all the viral contigs were combined for subsequent analysis.

### Viral Contigs Cluster and Quality Control

We applied CheckV v.1.0.1 (database v.1.4) to assess the quality of all viral sequences (Nayfach et al., 2021a). All viral contigs were clustered into species-level vOTUs based on 95% average nucleotide identity (ANI) and 85% alignment fraction (AF) using pairwise ANI calculations from the CheckV repository’s rapid genome clustering supporting code. All-vs-all local alignments were performed with blast+ package v.2.13.0, with the parameters (-max_target_seqs 10000). Pairwise ANI values were calculated by combining local alignments between sequence pairs using anicalc.py script. UCLUST-like clustering was carried out with the aniclust.py script using the recommended-parameters (95% ANI + 85% AF) from MIUVIG. The sequences of vOTUs resulting from clustering were extracted from the viral contigs, and their quality was re-evaluated using CheckV.

### Viral Taxonomy Annotation

Viral taxonomy identification was challenging due to the lack of a specific marker gene for viral sequence and the presence of a large amount of ‘dark matter’ in various environments. The new ICTV taxonomy was released in 2022 and abolished the previous large proportion of the order *Caudovirales* and the families *Myoviridae*, *Siphoviridae* and *Podoviridae* (Turner et al., 2023). To obtain more taxonomy information for vOTUs, we applied various software tools, including vConTACT2 v.0.11.3 (Bin Jang et al., 2019), geNomad v.1.2.0 (Camargo et al., 2023b), DemoVir script (https://github.com/feargalr/Demovir), CAT v.5.2.3 (von Meijenfeldt et al., 2019), and Kaiju v.1.9.0 (Menzel et al., 2016). All these methods used the new ICTV taxonomy database (v.214). We downloaded the reference sequence from NCBI according to the newest genebank-ID based on ICTV taxonomy and created new databases for each software tool following their respective manuals. If the protein sequences were needed, prodigal-gv v.2.9.0 was used to predict them (Hyatt et al., 2010). We also employed the blastn method against the IMG/VR4 database with the following parameters: -max_target_seqs 1000 -evalue 1e-5 -perc_identity 90 -qcov_hsp_perc 80 (Camargo et al., 2023a). The taxonomy of vOTUs was determined based on the results of various analysis methods in the following order: geNomad-blastn-vConTACT2-CAT-Kaiju-DemoVir.

### Functional Annotation

All protein-coding genes of vOTUs were predicted using prodigal-gv v.2.9.0 (Hyatt et al., 2010). Genes were annotated based on Dimond searches against protein databases including eggNOG 5.0 and VOGDB (http://vogdb.org) (Huerta-Cepas et al., 2018) using EggNOG-mapper with default parameters (Cantalapiedra et al., 2021). The structured antibiotic-resistance gene (SARG) database (Yin et al., 2022), BacMet database (Pal et al., 2013), mobileOG-db (Brown et al., 2022), VFDB(Liu et al., 2021a), and quorum sensing database (Wu et al., 2022b) were searched against with blastp using Diamond v.2.0.15.153 (Nayfach et al., 2021b) with specific parameters: --evalue1e-5 –query-cover 85 –id 60 (Buchfink et al., 2021).

### Virus Host Prediction

To predict the hosts of vOTUs, we aggregated data from several sources. The GTDB database (release 207_v2), containing a total of 317,542 genomes of bacteria and archaea, was used to serve as the host reference database (Chaumeil et al., 2022). For host predictions based on the Markov model, WIsH was performed against the GTDB database using criteria *P* value ≤ 0.05 (Galiez et al., 2017). In cases where the WIsH method failed, we utilized the blastn method against the IMG/VR4 database with previously predicted hosts for viral genomes in our vOTU database belonging to the same species (95% ANI + 85% AF) for cases where the WIsH method failed.

### Identifying Temperate Phages

We used BACPHLIP v.0.9.6 (Hockenberry & Wilke, 2021) and PhaTYP (Shang et al., 2022), which employed bidirectional encoder representations from transformer (BERT), to determine whether the vOTUs were likely to be temperate or virulent. However, since BACPHLIP’s result would indicate virulence if no protein domains of interest were found, the result for all vOTUs was heavily skewed towards virulent phage.

### Comparison to Other Viral Reference Databases

In this study, the vOTUs from the PVD were compared against four reference databases: IMG/VR v.4.0 (Camargo et al., 2023a), GVD v.2.0 (Gregory et al., 2020), MGV v.1.0 (Nayfach et al., 2021b), and GPD v.1.0 (Camarillo-Guerrero et al., 2021). To improve computational efficiency and facilitate cross-database comparison, all sequences were filtered to the vOTU species level and above 50% completeness according to the checkV results before clustering between different databases. The sequences were then combined and clustered using the supporting code in the checkV repository with blastn specific parameters: evalue 1e-5, max-target-seqs 10,000, query coverage 90, and identity 70. We extracted the VCs between different databases, and if the sequences clustered together, it indicated that the vOTU was shared between two databases, otherwise it meant that they were different. The comparison results were visualized using the UpSetR package (Conway et al., 2017).

### Phylogenetic Analyses

We generated phylogenetic trees for the vOTUs identified with taxonomy and host vertebrates, nucleocytoplasmic large DNA viruses (NCLDVs), and archaea with completeness above 50%. First, we performed multiple sequence alignment for each type of vOTU sequence using MAFFT v.7.515 with the “—auto” model (Katoh et al., 2002) for each type of vOTU sequence. Second, we inferred a concatenated nucleic acid sequence phylogeny from the multiple sequence alignment using FastTree v2.1.11 with the parameters “-nt-gtr” (Price et al., 2009). Finally, the tree was midpoint rooted and visualized using iToL (Letunic & Bork, 2021).

## Results

### The DNA virus from the pig gut microbiome

In this study, our objective was to create a comprehensive pig virome database (PVD) of the gut utilizing next-generation sequencing of metagenomic samples and the development of metav, a virus detection pipeline that integrates established methods and signatures such as VirSorter2 and DeepVirFinder for viral sequence identification (Guo et al., 2021a; Ren et al., 2020). Our findings indicated that the choice of assembler, specifically Megahit versus metaSPAdes, had a minimal effect on the recovery of viruses, which was consistent with prior research (Nayfach et al., 2021b). We used the Megahit assembler and selected -k-list 39, 59, and 79 after testing different kmer lists based on computational efficiency. We applied metav to 4,650 faecal and gut content samples from various regions worldwide, including the USA, China, Europe, and Australia, resulting in the identification of 19,577,760 unique, single-contig viral genomes exceeding 1.5 kb in length, which allowed us to comprehensively capture DNA viruses in the pig gut (Fig. 1a and Extended Data Fig. 1). By clustering viral contigs using 95% ANI and 85% AF, we identified 3,219,423 viral operational taxonomic units (vOTUs) at the species level (Fig. 1a). In addition, we evaluated vOTU genomes quality using CheckV and found 48,299 vOTU genomes (corresponding to 5,566,804 viral contig sequences) that were at least 50% complete (Nayfach et al., 2021a), including 12,515 complete vOTU genomes (corresponding to 2,099,394 viral contig sequences) (Fig. 1b and c, Supplementary Table 1). Notably, although only 1.50% of total vOTU genomes were above 50%, they corresponded to 28.43% of contig sequences with >50% completeness to total contig genomes, suggesting that small genome fragments harbour greater viral diversity (Fig. 1d). Moreover, the size of most vOTU genomes with completeness exceeding 50% was above 10 kb, indicating their potential to provide more reliable results (Khan Mirzaei et al., 2021; Nayfach et al., 2021b).

**Fig. 1.**
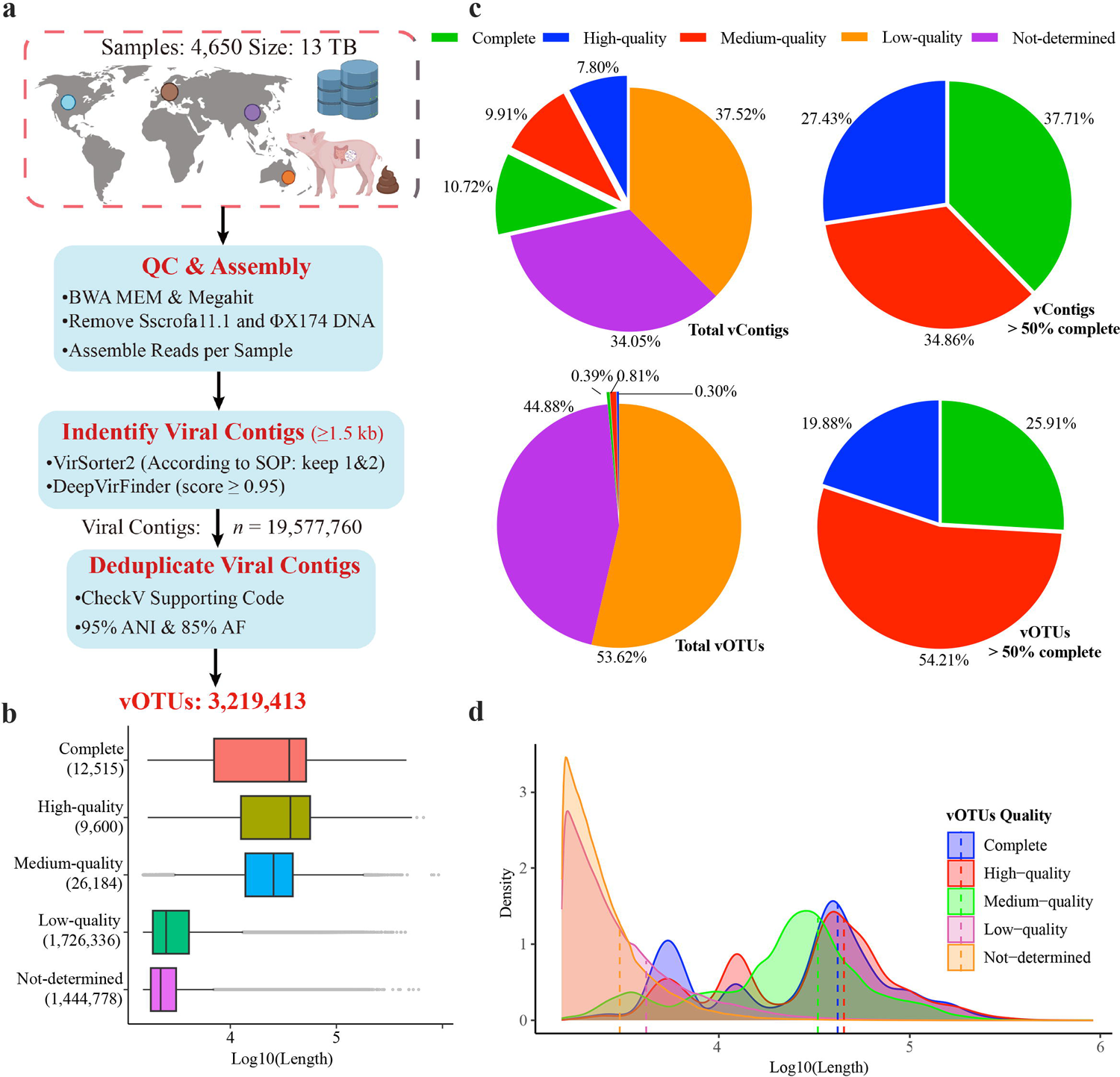
Summary of Viral Genomes Recovered from Pig Gut Metagenomes. **a**, Overview of the PVD Database viral discovery pipeline. **b**, Distribution of estimated genome completeness for viral operational taxonomic units (vOTUs) in the PVD database grouped into quality tiers: Complete (n = 12,515), High Quality (n = 9,600), Medium Quality (n = 26,184), Low Quality (n = 1,726,336), and Not Determined (n = 1,444,778). **c**, Proportion of viral contigs and vOTU genomes based on quality tiers. **d**, Distribution of density for different vOTUs quality.

### Taxonomic Annotation

Utilizing the new ICTV classification, which abolished the paraphyletic morphological families *Podoviridae*, *Siphoviridae*, and *Myoviridae*, as well as the order *Caudovirales*, and the families and order were replaced them with the class *Caudoviricetes* to categorize all tailed bacterial and archaeal viruses featuring icosahedral capsids and a double-stranded DNA genome (Turner et al., 2023). After using the new ICTV classification, the majority of vOTUs (65.36%) were identified as *Caudoviricetes* (Fig. 2a, Extended Data Fig. 1, and Supplementary Table 2). However, despite using various techniques to capture the taxonomy comprehensively, only 18.43% of vOTU genomes could be annotated at the family level using the new ICTV database (Fig. 2b and Extended Data Fig. 2). These findings suggested significant gaps in knowledge regarding pig gut virus taxonomy, which is also a significant challenge in analysing virome from different environments (Breitbart et al., 2003; Liang & Bushman, 2021; Nayfach et al., 2021b). Our taxonomy classification results combined with the new ICTV list showed that bacteria were the most common hosts, accounting for 95% (n=563,390) of the total taxonomy identification vOTU genomes (Fig. 2b). At the family level, dominant vOTUs infecting bacteria included *Peduoviridae* (n=368,019), *Herelleviridae* (n=52,947), *Steigviridae* (n=34,953), *Suoliviridae* (n=17,276), and *Straboviridae* (n=12,993). Archaeal hosts were predominantly infected by *Vertoviridae* (n=4,382), *Anaerodiviridae* (n=879), *Halomagnusviridae* (n=602), *Hafunaviridae* (n=404), *Saparoviridae* (n=261), and *Lipothrixviridae* (n=222). Among vertebrates, *Poxviridae* (n=2,004), *Iridoviridae* (n=1,102), *Herpesviridae* (n=308), *Smacoviridae* (n=289), *Circoviridae* (n=247), and *Retroviridae* (n=200) were the dominant taxonomies. We also identified several vOTUs belonging to the nucleocytoplasmic large DNA viruses (NCLDVs), including *Phycodnaviridae* (n=6,646), *Mimiviridae* (n=6,306), *Marseilleviridae* (n=983), and *Ascoviridae* (n=279), with *Poxviridae* and *Iridoviridae* capable of infecting vertebrates (Fig. 2b).

**Fig. 2.**
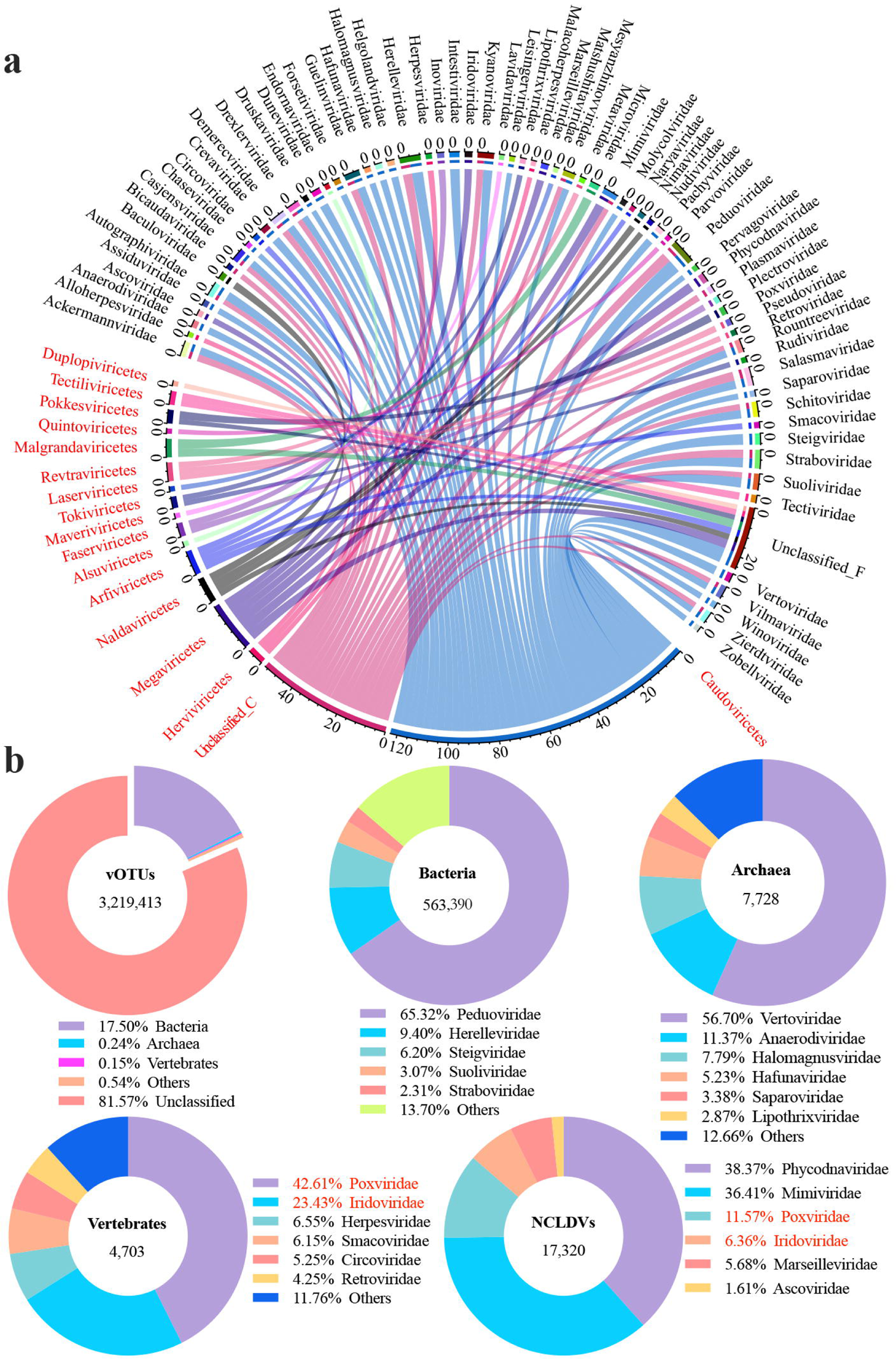
The taxonomy and proportion of host assignment of vOTUs. **a**, Circle chart that depicts the vOTUs taxonomy at both the class and family levels. (The names of classes are shown in red font, and those of families in black font). **b**, Proportion of vOTUs and viral genomes that belong to various hosts, as well as nucleocytoplasmic large DNA viruses (NCLDVs) with identified taxonomy. The red font highlights the percentage of families shared by vertebrate hosts and NCLDVs.

### Host Prediction and Temperate Identification

Accurately predicting virus host is a critical step towards understanding virus-host interactions and utilizing them to manipulate the gut microbiota ecosystem or design innovative phage tools (Kauffman et al., 2022; Khan Mirzaei & Deng, 2022). In this study, we employed the WIsH tool to target the GTDB database and predict the hosts of vOTU genomes. As expected, Firmicutes and Bacteroidetes, the most abundant bacterial phyla in the gut microbiome, were found to be common hosts for viruses at the phylum level (Fig. 3a and b). At the genus level, *Clostridium* (belonging to Firmicutes) and *Prevotella* (belonging to Bacteroidetes) were the most commonly assigned hosts (Fig. 3a). Additionally, some viruses were predicted to inhabit *Ruminococcus, Ruminiclostridium, Blautia, Faecalibacterium,* and *Eubacterium*, which are known to play important roles in pig productivity and feed efficiency (Bergamaschi et al., 2020; Ramayo-Caldas et al., 2016). The ratio of virulent and temperate phages varied among the different hosts, with most phages predicted to be virulent (Fig. 3a).

**Fig. 3.**
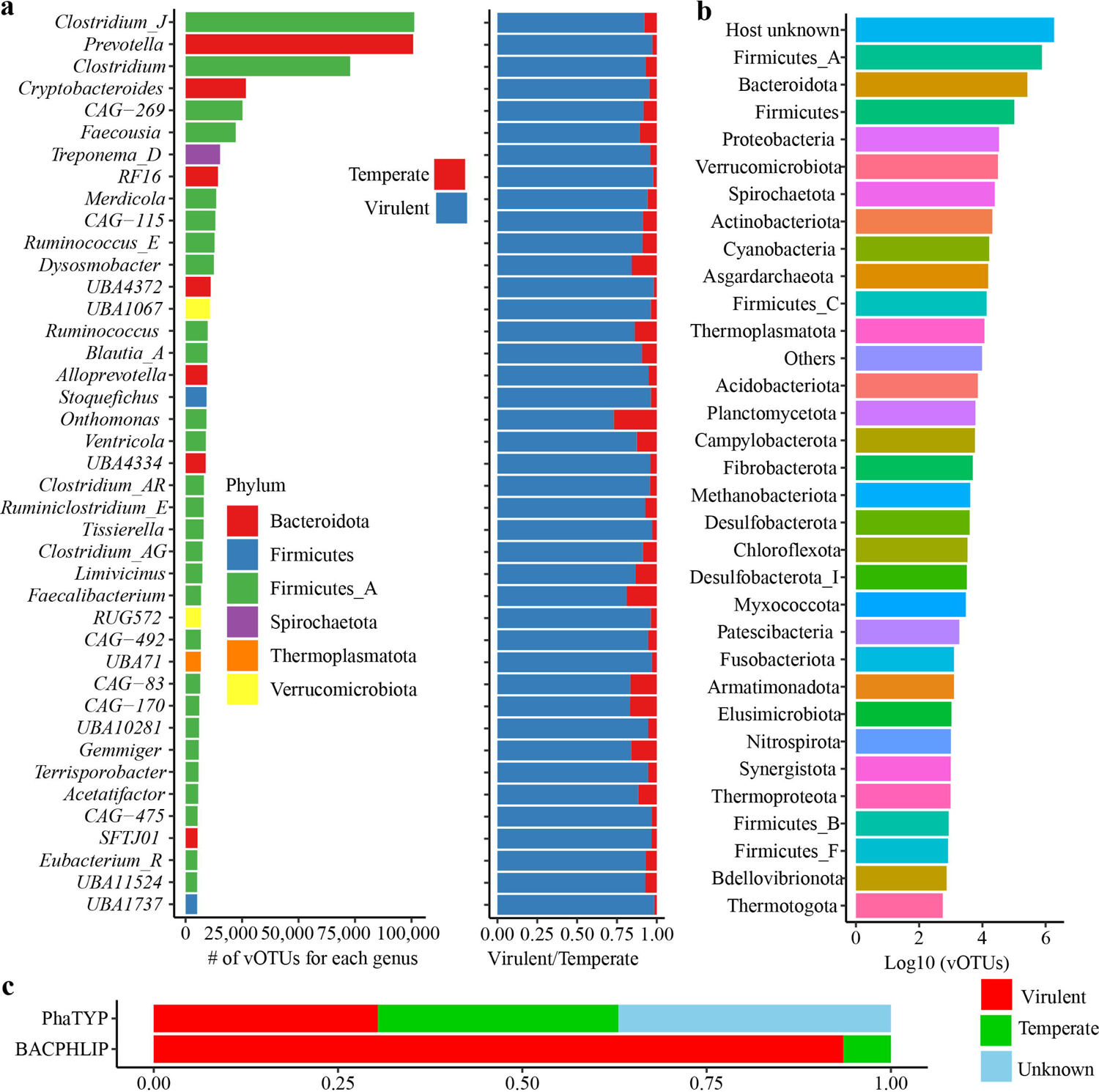
Host prediction and temperate identification of vOTUs. **a**, Number of vOTUs and the ratio of virulent and temperate phages for each predicted host at the genus level. **b**, Number of vOTUs for each predicted host at the phylum level. **c**, The ration of virulent and temperate phages for total vOTUs indentified by PhaTYP and BACPHLIP.

We used BACPHLIP analysis to classify the majority of vOTU genomes (93.53%) as virulent (Fig. 3c) (Hockenberry & Wilke, 2021). However, previous studies have shown that incompleteness is not the main factor affecting the detection of virulent viruses, and it is not uncommon to recover genome sequences of lytic viruses from stool metagenomes (Nayfach et al., 2021b). Therefore, we utilized PhaTYP, a python library for bacteriophage prediction with a BERT-based model, to confirm the lifestyle of vOTUs in our study. The results showed that temperate, virulent, and unknown categories accounted for 32.62%, 30.41%, and 36.97% of all predictions, respectively. However, other studies have reported that lytic viruses occupy over 90% of the total viruses in the ocean and human gut environments (Luo et al., 2022; Nishijima et al., 2022). These discrepancies suggest that further research is needed to determine the reasons for differences in predicting the lifestyles of viruses.

### Newly Established and Expanded Pig Gut Viral Diversity

To ensure comparability and consistency with established quality standards for genomic comparisons (Bowers et al., 2017; Roux et al., 2019), we focused on 48,299 vOTU genomes with more than 50% completeness. We performed checkV on the IMG/VR4, MGV, GVD, and GPD datasets to remove low-quality sequences, retaining vOTU genome fragments with more than 50% completeness. Following previous studies (Camarillo-Guerrero et al., 2021; Nayfach et al., 2021b), we combined all vOTU genomes >50% from these four datasets with those from the PVD database and clustered them at the species level using a query coverage of 0.9 and identity of 0.7. Our results showed that the PVD database significantly enhanced the diversity of viral genomes from pig gut microbiota metagenomic samples. Of the 47,650 species-level vOTUs in the PVD database, 34,952 (92.83%) did not cluster with any vOTU genomes from the other datasets (Fig. 4). In contrast, the IMG/VR4 database contained approximately 189, 40, 35, and 32 times higher numbers of viral genomes than the GVD, PVD, MGV, and GPD databases, respectively. Of the 1,504,202 vOTU genomes from the IMG/VR4, 1,455,627 (96.77%) were unique compared to the other databases (Fig. 4). However, human gut metagenome samples used to construct the GVD, MGV, and GPD databases represented a higher proportion of viral genomes in the IMG/VR4 database than pig gut viruses. Specifically, GVD, MGV, and GPD had 4,712 (58.79%), 43,464 (95.19%), and 25,377 (53.56%) sequences from human gut metagenomes, respectively, while GPD, GVD, and MGV had 20,695 (43.68%), 3,247 (40.51%), and 1,910 (4.39%) unique sequences compared to the other databases, respectively (Fig. 4). Our findings highlighted the importance of using comprehensive and high-quality databases to advance our understanding of viral diversity in different environments. The improved detection of viral reads in whole metagenomes and expanded coverage of virus‒host connections in the PVD database make it particularly useful for studying pig gut viruses.

**Fig. 4.**
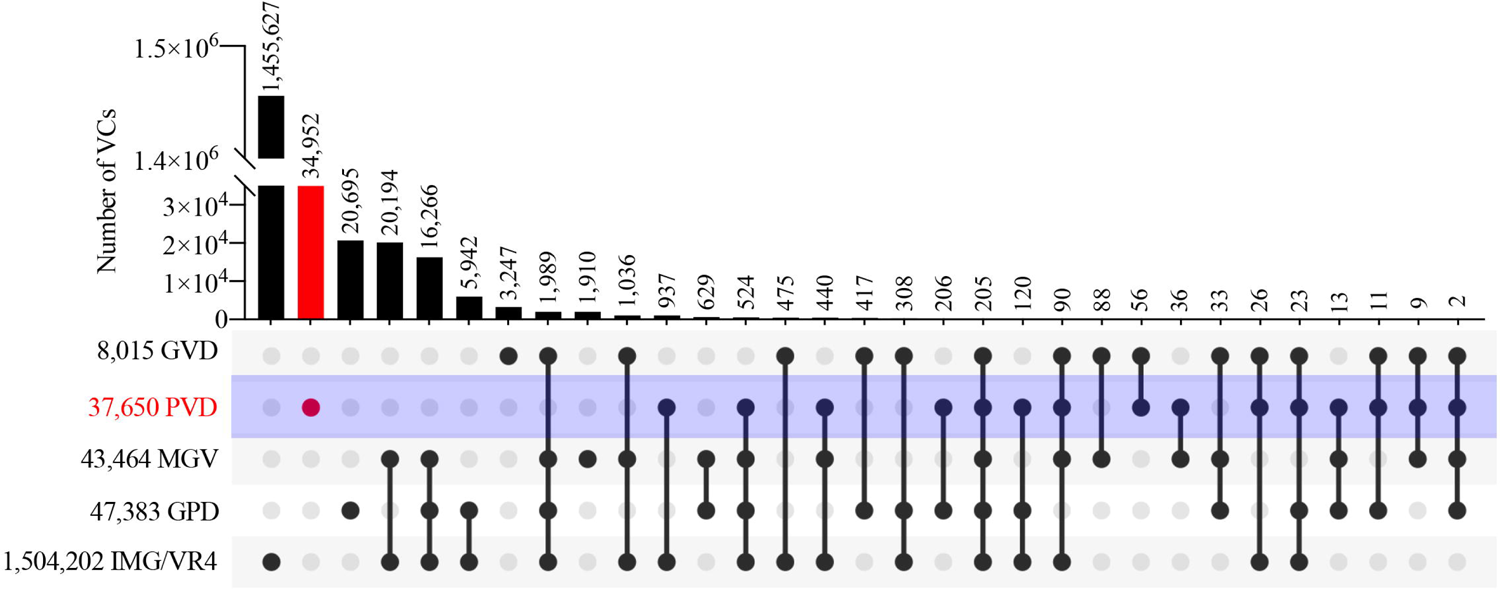
vOTUs clustering and comparison with four existing datasets. The vOTU genomes from the PVD (n = 37,650) catalogue were compared with other virus genomes >50% completeness from four databases: IMG/VR4 (n = 1,504,202), MGV (n = 43,464), GPD (n = 47,383), and GVD (n = 8,015) at species level with parameters query coverage 90, and identity 70.

### Phylogenomic Analyses of Viruses

Viruses that infect vertebrates can cause serious diseases in pigs, leading to significant economic losses for the swine industry. For example, porcine circovirus can cause porcine circovirus-associated disease (PCVAD) (Díaz et al., 2021; Klangprapan et al., 2021). In our metagenomic samples, we did not detect *Asfarviridae*, a large encapsulated double-stranded DNA virus (Chen et al., 2019), possibly because the samples were collected from healthy or noninfected pigs. To explore the diversity of pig gut viruses, we constructed a species-level phylogenetic tree based on the 597 genomes with >50% completeness from PVD (Fig. 5). The majority of viruses from the PVD represented new lineages across the tree. *Smacoviridae*, *Circoviridae*, and *Redondoviridae* were the most diverse families found, primarily due to the broad phylogenetic distribution of the vOTUs belonging to these groups, which are circular replication-associated protein (Rep)-encoding single-stranded DNA viruses (Abbas et al., 2019; Liu et al., 2021b; Varsani & Krupovic, 2018). Notably, *Redondoviridae* is the second most prevalent eukaryotic DNA virus family and was first reported and named using metagenomics identification methods, which were associated with human diseases (Abbas et al., 2019; Varsani & Krupovic, 2018). Moreover, we found porcine circovirus-like viruses that have been associated with porcine diarrhoeal disease (Liu et al., 2021b). Large and giant viruses belong to a group of double-stranded DNA viruses known as nucleocytoplasmic large DNA viruses (NCLDVs), which constitute the viral phylum *Nucleocytoviricota* (Koonin et al., 2020; Schulz et al., 2022; Schulz et al., 2020). These viruses infect a wide range of eukaryotic hosts, and the viruses have genome sizes ranging from 70 kb to up to 2.5 Mb (Schulz et al., 2022; Sun et al., 2020). In our current study, two genomes belonging to *Mimiviridae* had more than 100 kb (Extended Data Fig. 3). Additionally, we found 74 archaeal genomes with >50% completeness that were dominated by *Anaerodiviridae* and *Vertoviridae*, which was different from the archaeal virome in the human gut (Extended Data Fig. 4) (Li et al., 2022b). Compared with recently published collections of viruses from diverse environments and the human gut (Al-Shayeb et al., 2020; Nayfach et al., 2021b), our analysis of the PVD resulted in a substantial expansion of pig gut viral diversity. The PVD provides valuable references for virus research, as well as for the prevention, control, pathogenesis, and epidemiology of these viruses.

**Fig. 5.**
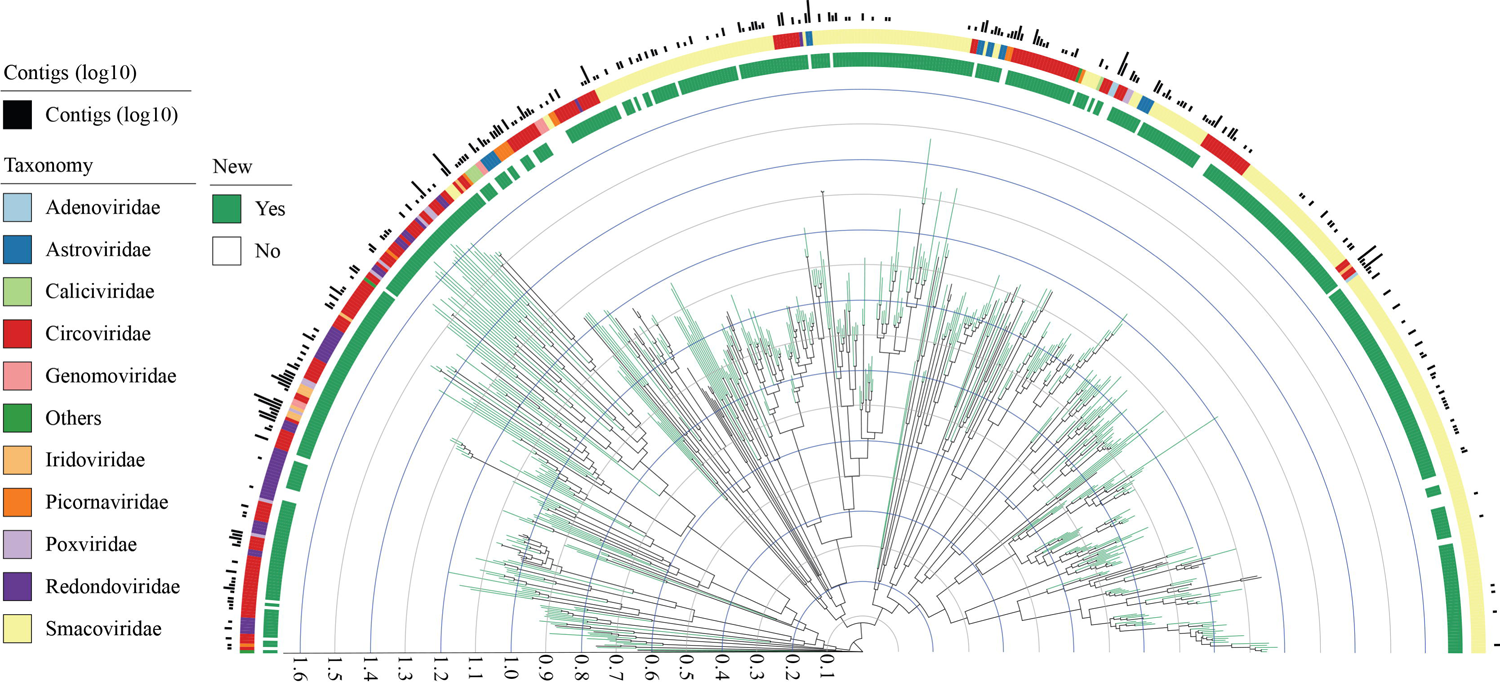
Phylogenomics of vOTUs habited vertebrates. A phylogenetic tree was constructed from 597 species-level derived from the PVD. Tree was plotted using iTOL. Branch colour indicates whether a lineage is represented by previously polished databases (IMG/VR4, MGV, GPD, and GVD) (black) or is unique to the PVD database (green). Outer rings display different results for each vOTU.

### Functional Capacity of the Gut Virome

To investigate the potential roles of the pig gut virome, we identified 18,161,503 protein-coding genes across 3,219,413 vOTU genomes from our current study. We then compared these genes to databases such as SARG, BacMet, mobileOG-db, VFDB, and the quorum sensing database using Diamond’s blastp tool, with parameters set to --evalue1e-5, --query-cover 85, and --id 60 (Fig. 6a). Our analysis revealed that phages, which constitute the largest proportion of the gut virome, could potentially facilitate the spread of antibiotic resistance genes (ARGs) through conduction (Calero-Caceres et al., 2019; McInnes et al., 2020; Sun et al., 2021). The pig faecal virome exhibits the highest relative abundance of antibiotic resistance genes (ARGs) compared to other environments (Calero-Caceres et al., 2019). The virome also carries antibiotic resistance genes (ARGs) in other environments, such as soil, rivers, and faeces (Chen et al., 2021c; Karkman et al., 2019; Moon et al., 2020). We identified 4,564 protein-coding genes targeting the SARG database for antibiotic resistance genes (ARGs), accounting for only 0.25‰ of the total protein genes. The top five categories were polymyxin (n=1,803), macrolide-lincosamide-streptogramin (n=1,200), aminoglycoside (n=335), vancomycin (n=329), and tetracycline (n=291) (Fig. 6a). However, we observed that the parameters of query cover and identity had a significant impact on the number of ARGs detected. Various studies have used different parameters to detect ARGs carried by viruses, often below the 60% identity threshold used in our study and recommended by Yin et al. (2022). This could lead to inaccuracies and overestimations in the reported number of ARGs (Chen et al., 2021c; Moon et al., 2020). For example, by setting the parameters of –query-cover 60 and –id 40 in blastp, we obtained 34,993 hits (0.19%), which is 7 times higher than the previous setting. To compare results across different studies and establish more appropriate parameters for detecting virus-carried ARGs, comprehensive evaluations of parameters should be conducted in the future (Billaud et al., 2021; Enault et al., 2017). We also explored the distribution patterns of virulence factor genes (VFGs) carried by the virome (Fig. 6a). The results showed that VFGs were mainly induced by the immune modulation (n=15,698), adherence (3,731), and stress survival (1,243), meaning that the virome also plays very important roles in the spread of VFGs to pathogenetic and antimicrobial-resistant bacteria (Liang et al., 2020).

**Fig. 6.**
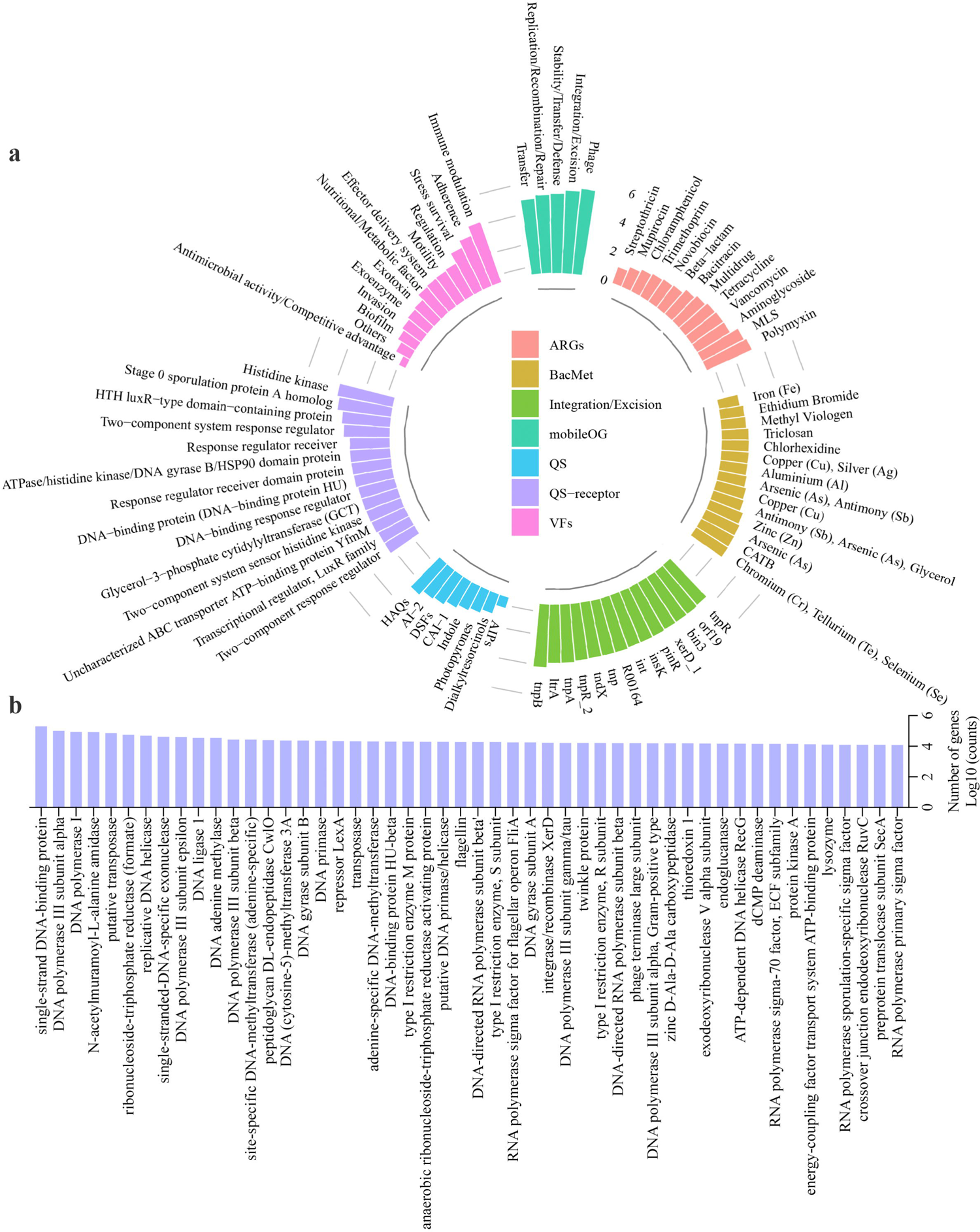
Functional landscape of pig gut virome. **a**, Protein-coding viral genes were identified across all vOTUs from PVD and compared with protein profiles from SARG, BacMet, mobileOG-db, VFDB, and quorum sensing databases with blastp using Diamond with specific parameters: --evalue1e-5 –query-cover 85 –id 60. **b**, Functional annotations for the largest 50 protein clusters.

We utilized the recently developed mobileOG-db to detect mobile genetic elements (MGEs) in pig gut vOTUs (Fig. 6a). This database contains more than 700,000 deduplicated sequences specifically designed for MGE detection (Brown et al., 2022). A total of 1,145,163 protein-coding genes (6.31% of total protein genes) were found and encompassed five major categories: phage (n=439,812), integration/excision (n=287,492), stability/transfer/defense (n=175,226), replication/recombination/repair (n=150,497), and transfer (n=92,136) (Fi. 6a). Our subsequent analysis focused on the minor categories of integration/excision, which may contribute to the spread of ARGs. Among these, tnpB (n=14,287) and tnpA (n=6,892) were the most abundant and are known to play an important role in the transposition of ARGs (Karvelis et al., 2021). Our findings indicate that the virome of pig gut also harbours integron genes (int/int1, n=4,745), which are crucial for the spreading of antibiotic resistance genes (ARGs) (Zhang et al., 2020). Moreover, exposure to metals and biocides may co-select for resistance in the microbiome (Li et al., 2022c). Notably, we detected a small number of BacMet resistance protein genes (n=2,185) in the pig gut vOTUs (Fig. 6a). The majority of metal resistance genes in the pig gut virome belonged to chromium, tellurium, and tselenium (Cr/Te/Se) (n=308), arsenic (As) (n=276), zinc (Zn) (n=216), and copper (Cu) (n=105). Cu and Zn are extensively used as feed additives for pigs and have a significant impact on the dissemination of antibiotic resistance genes (ARGs) in manure and soil (Li et al., 2022a). Taken together, our results suggested that the pig gut virome played a crucial role as a reservoir and source of ARGs dissemination. However, further studies are needed to investigate the factors and mechanisms underlying the spread of ARGs by gut viruses.

While the quorum sensing of bacteria has received considerable attention (Whiteley et al., 2017), our understanding of gut virus quorum sensing remains limited. To address this gap, we examined vOTU proteins against the Quorum Sensing of Human Gut Microbes (QSHGM) database (Wu et al., 2022b). We identified only 1,164 protein genes of pig gut vOTUs that utilized quorum sensing languages, with the top three languages being HAQs, AI-2, and DSFs (Fig. 6a). These results were consistent with those reported for the human gut microbiome (Wu et al., 2022b). However, the pig gut virome had a higher number of quorum sensing receptors (n=20,627), with histidine kinase, stage 0 sporulation protein A homologue (Spo0A), LuxR, and a two-component system response regulator being the dominant receptors. These receptors may be crucial for maintaining the balance of the gut microbiota ecosystem through communication between viruses and hosts (Butala & Dragoš, 2022; Leeks et al., 2021). Nevertheless, further research is needed to investigate quorum sensing systems and establish a dedicated database for pig gut microbiota.

To explore the functional potential of the pig gut virome, we used eggNOG-mapper v.2 to compare against eggNOG 5.0 using default parameters (Cantalapiedra et al., 2021; Huerta-Cepas et al., 2018). Overall, 51% (9,326,889/18,161,503) of vOTU genes had matched to the database (Fig. 6b). However, a striking 91% of these genes were not assigned any biological function, highlighting our limited understanding of the pig gut virome. Among the matched genes, the largest protein clusters were single-stranded DNA-binding protein (n=201,294), followed by other typical viral functions such as DNA polymerase and replicative DNA helicase (Fig. 6b). With respect to potential metabolic functions, we detected a total of 543,051 genes, with the top five categories being carbohydrate metabolism (n=123,948), nucleotide metabolism (n=105,153), amino acid metabolism (n=80,577), metabolism of cofactors and vitamins (n=55,590), and energy metabolism (n=51,182) (Extended Data Fig. 5).

## Discussion

In the current study, we conducted a comprehensive analysis of pig gut viral genomes by mining publicly available metagenomic samples (n=4,650) using our integrated pipeline called metav (www.github….). The software tools in this pipeline were optimized to improve the speed and accuracy of metagenomic analysis and enable comparisons between different microbial ecosystems. We identified 19,577,760 draft-quality viral genomes, representing an estimated 3,219,413 species-level vOTUs. After applying CheckV, the PVD database was established, which includes 5,566,804 viral contigs with more than 50% completeness, forming 48,299 species-level vOTUs. This is equivalent to the MGV (n=54,118) and GPD (n=46,480) human gut virome databases (Camarillo-Guerrero et al., 2021; Nayfach et al., 2021b). However, a pair-to-pair clustered comparison between IMG/VR4, GPD, MGV, GVD, and PVD showed that PVD still has 34,952 (92.83%) unique vOTUs, making it the largest resource containing extensive pig gut viral genomes to the best of our knowledge. Although this study focused on DNA viruses, other studies have explored RNA viruses (Chen et al., 2022; Dominguez-Huerta et al., 2022; Edgar et al., 2022; Neri et al., 2022; Shkoporov et al., 2022b; Zayed et al., 2022). Thus, future investigations could utilize metatranscriptomics data from pig gut microbiota samples to investigate RNA viromes.

To rapidly recover viral genomes from metagenomics and metatranscriptomics data, ICTV has updated its taxonomy list, which has brought about significant changes compared to the previous version (Turner et al., 2023). However, this modification has made it difficult to compare the taxonomic results of this study with those of previous studies. To address this issue, it would be useful, in the future, to integrate several large-scale viral genome databases and create a unified, updated, and standardized catalogue that is compatible with the new ICTV taxonomy. Different taxonomy software can produce conflicting results, and some may not support the new referenced taxonomy classification (Extended Data Fig. 2 and Supplementary Table 2). The classic software vConTACT2, which clusters and provides taxonomic context for viral metagenomic sequencing data, is no longer being actively developed and is time-consuming for large database (Bin Jang et al., 2019). In contrast, geNomad is significantly faster than similar tools and can process large datasets, accurately assigning identified viruses to the latest ICTV taxonomy release (Camargo et al., 2023b). Another promising program, PhaGCN, uses a novel semisupervised learning model for taxonomic classification of phage contigs (Shang et al., 2021). However, it is not supported for contigs below 2 kb. Therefore, it is essential to establish a unified and standardized taxonomy classification pipeline or platform in the future (Bolduc et al., 2021).

The results of our study provided significant insights into the composition and function of the pig gut virome. Our analysis uncovered a diverse and complex pig gut virome, consisting of numerous viral operational taxonomic units (vOTUs). In addition, we have identified several potentially novel viral species and found evidence of multiple viral species coinfecting the pig gut. These discoveries highlight the importance of continued research in this field, as they could have significant implications for our understanding of viral evolution, pathogenesis, and host‒pathogen interactions. Moreover, our findings suggest that coinfection with multiple viral species is common in the pig gut, which has important implications for future studies investigating the impact of the pig gut virome on pig health and productivity, as well as the potential for viral transmission to other animals and humans. Given the lessons learned from the COVID-19 pandemic, such as the potential for emerging infectious diseases to adversely affect human health adversely, there is a need for increased awareness of zoonoses (Banerjee & van der Heijden, 2022; Ko et al., 2022). Therefore, we propose a “One Health” framework that emphasizes the importance of studying the “ecosystem-animal-livestock-human” pathogen system using metagenomic technology. This approach should include combining rich metadata, such as climate change data, to reveal the sources, hosts, transmission, and evolutionary mechanisms of known and unknown pathogens (Carlson et al., 2022). In summary, our study lays the groundwork for further research into the pig gut virome. Our identification of new viral species and evidence of coinfection highlight the intricacies of gut ecosystems and emphasize the importance of further investigation in this area. Our findings form the basis for a more comprehensive understanding of viral and microbial ecology in the pig gut and are expected to facilitate the monitoring and maintenance of pig health.

## Acknowledgements

This study was supported by the International Collaboration 111 Programme (BP0719040). We would like to express our gratitude to the Supercomputing Center of Lanzhou University for their valuable support in the computation works.

## Data availability

Access to the full vOTUs genomes from PVD is provided at https://doi.org/10.6084/m9.figshare.22671076.v1. Any further requests for data should be directed to the corresponding authors.

## Code availability

Supporting code is provided at https://github.com/mijiandui/metav.

## Author Contributions

J.M. designed the study, plotted the figures, and wrote the manuscript. J.M., Y.L., and Y.Z. collected the metagenomic data. Y. Y. provided suggestions. J.M., and C.M. performed the metagenomic analysis. X.J., R.L., and H.Z. contributed to the scientific discussion and preparation of the manuscript.

---

Extended Data Fig. 1 Geographic distribution of metagenome sampling sites involved in current study. The map of the sampling locations was created in ArcGIS v10.5. For each dot, the circle size reflects the number of metagenomic samples.

Extended Data Fig. 2 Taxonomy of vOTUs and the status of classification identified by different software. **a**, Number of vOTUs with bacteria as host according to ICTV list information at family level. **b**, Number of vOTUs with archaea as host according to ICTV list information at family level. **c**, Number of vOTUs with vertebrates as host according to ICTV list information at family level. **d**, Proportion and number of vOTUs identified by different software (Cat, Demovir, Kaiju, Phagcn Genomad, and Blastn) at family level, 1-means no conflict taxonomy result identified by different software. 2-meas conflict result with two software. 3-meas conflict result with three software. 4-meas conflict result with four software. **e**, Proportion and number of vOTUs with no conflict taxonomy result identified by different software at family level. **f**, Proportion and number of vOTUs identified by different software (Cat, Demovir, Kaiju, Phagcn Genomad, and Blastn) at class_family level, 1-4 were same means as show in d, 5-meas conflict result with five software. **g**, Proportion and number of vOTUs with no conflict taxonomy result identified by different software at class_family level. **g**, Proportion and number of vOTUs taxonomy identified by different software at class level.

Extended Data Fig. 3 Phylogenomics of vOTUs habited archaea. A phylogenetic tree was constructed from 74 species-level derived from the PVD. Tree was plotted using iTOL. Branch colour indicates whether a lineage is represented by previously polished databases (IMG/VR4, MGV, GPD, and GVD) (black) or is unique to the PVD database (green). Outer rings display different results for each vOTU.

Extended Data Fig. 4 Phylogenomics of vOTUs belong to nucleocytoplasmic large DNA viruses (NCLDVs). A phylogenetic tree was constructed from 37 species-level derived from the PVD. Tree was plotted using iTOL. Branch colour indicates whether a lineage is represented by previously polished databases (IMG/VR4, MGV, GPD, and GVD) (black) or is unique to the PVD database (green). Outer rings display different results for each vOTU.

Extended Data Fig. 5 Proportion of the function of vOTUs category as metabolism.

Supplementary Table 1 The quality of vOTUs checked by checkV. https://doi.org/10.6084/m9.figshare.22670755.v1.

Supplementary Table 2 The taxonomy classification with different software. https://doi.org/10.6084/m9.figshare.22671028.v1.

