## Supplementary figures and images for "Massive expansion of the pig gut virome based on global metagenomic mining"

### Extended Data Fig. 1

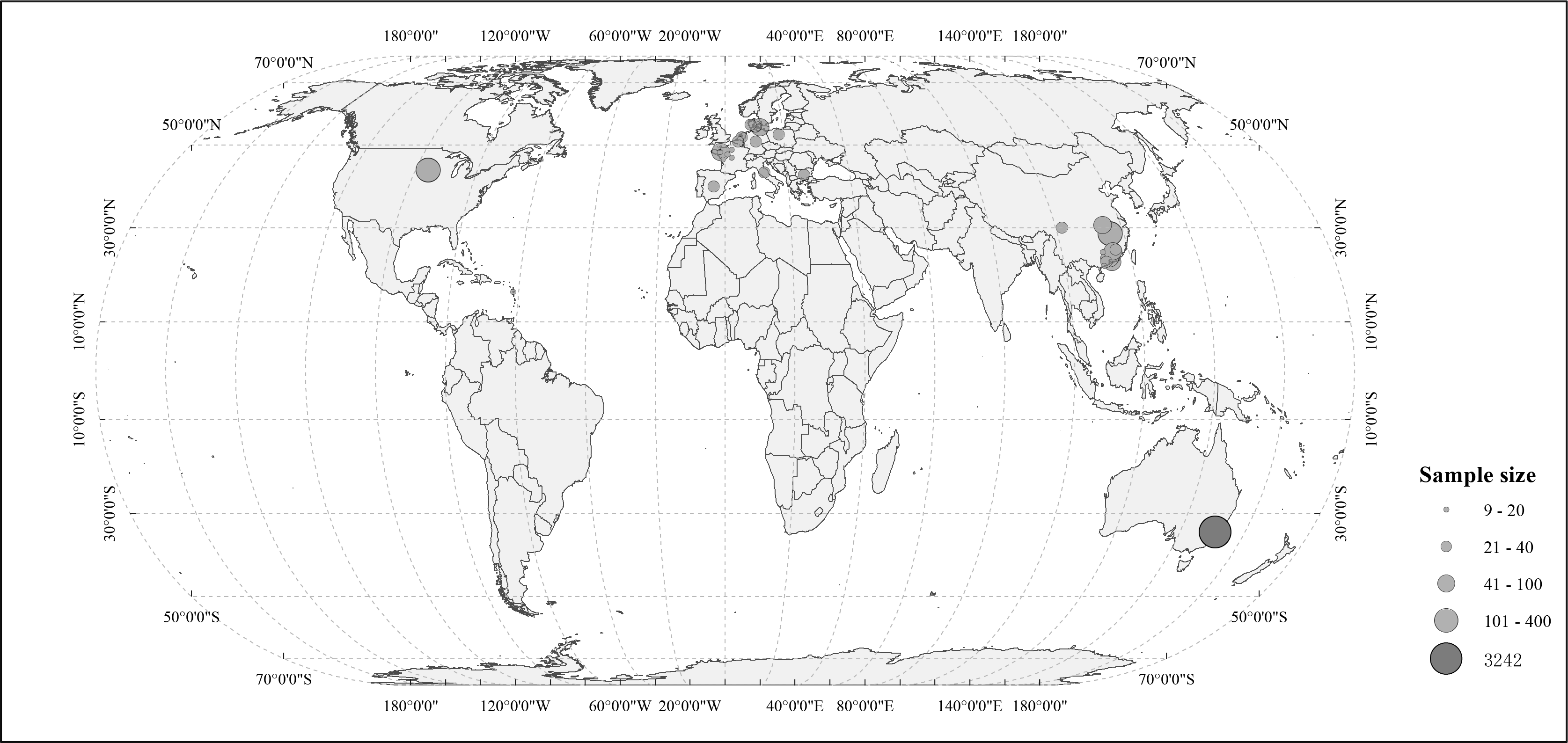

### Extended Data Fig. 2

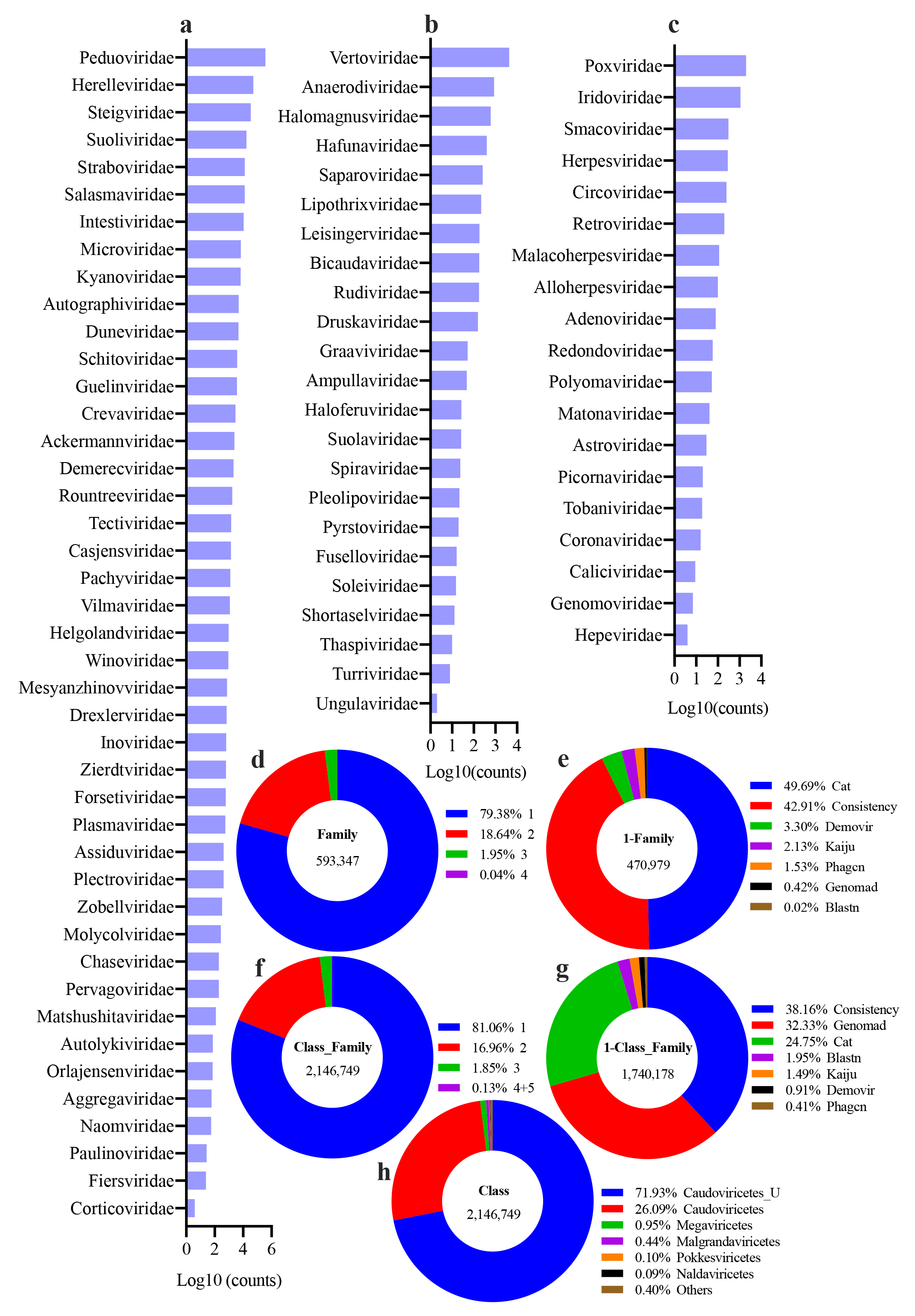

### Extended Data Fig. 3

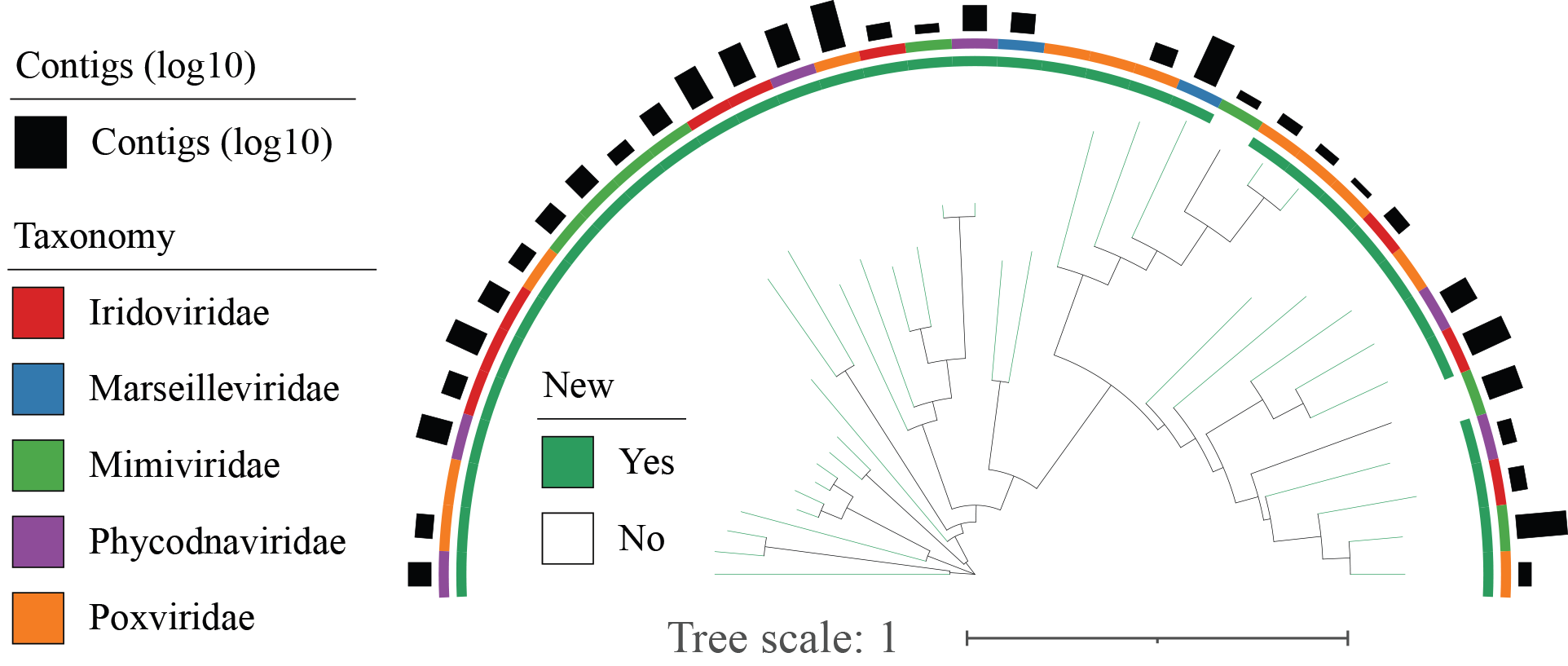

### Extended Data Fig. 4

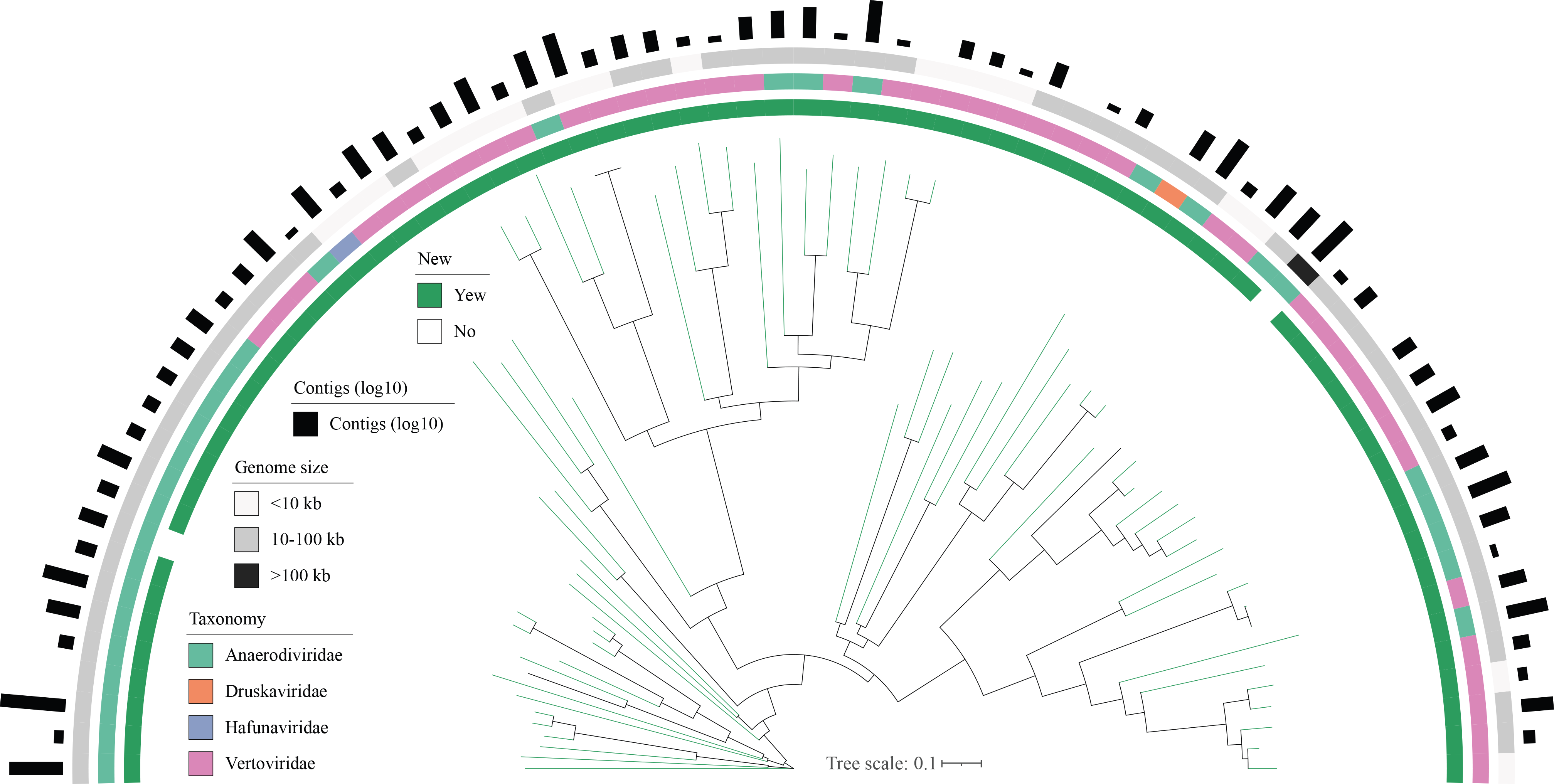

### Extended Data Fig. 5

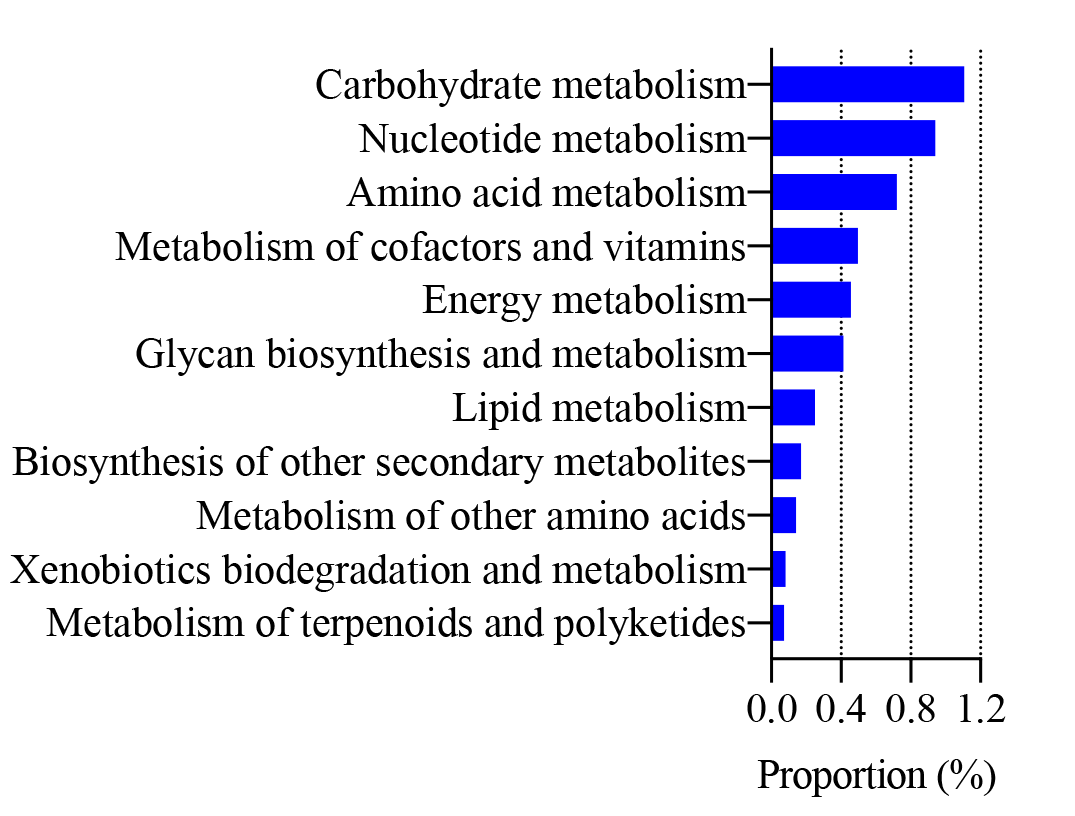
